# A systematic and meta-analytic review of non-verbal auditory memory in the brain

**DOI:** 10.1101/2024.11.11.622965

**Authors:** Fulvia Francesca Campo, Francesco Carlomagno, Betül Yılmaz, Luis Quaranta, Giulio Carraturo, Davide Rivolta, Elvira Brattico

## Abstract

While sounds are essential for human development, existing research primarily emphasizes visual and spatial memory and speech in relation to auditory processes, with a lack of a cohesive understanding of memory mechanisms for non-verbal sounds. This systematic review and coordinate-based meta-analysis of neuroimaging findings aims to comprehensively organize the literature for identifying the key brain mechanisms involved in short-term memory, working memory, and long-term memory for non-verbal auditory information. Additionally, we aimed to identify whether and how individual differences in neural memory processes related to auditory expertise (musicianship), aging or auditory impairments such as amusia are explored in the literature. Our review included ninety studies meeting the selection criteria, with only thirteen studies containing brain coordinates could be included in the meta-analysis. The coordinate-based meta-analysis identified a frontal hub for non-verbal auditory memory encompassing the medial frontal gyrus, cingulate gyrus and superior frontal gyrus.

## 1. Introduction

Sounds are crucial for human development, continuously influencing neural processes through neuroplasticity^1^. Despite this, current literature predominantly focuses on visual and spatial memory, as well as speech in relation to audition, while a unified perspective on memory processes for non-verbal sounds is lacking^1^. This review aims at organizing the neuroimaging literature on memory for non-verbal sounds.

Environmental sounds and noise are omnipresent in the auditory scene, influencing humans in both positive and negative ways. Pleasant sounds can enhance parasympathetic activity and lower cortisol, while unpleasant sounds trigger sympathetic responses and activate the amygdala^2–5^. Environmental sounds are non-human-made, whereas urban noise, often generated unintentionally by human activity, forms a chaotic auditory scene. Despite their randomness, our auditory system is programmed to recognize these sounds as “auditory objects” ^6^, organizing them preattentively (by segregation and integration, following Gestalt principles^7^) and attentively associate them with specific object categories^8^. Music, as a form of intentional, human-made sound, is one of the most complex non-verbal auditory stimuli, governed by culturally dependent structural principles^9,10^. Its comprehension requires higher-order cognitive processes like memory and attention from both listeners and musicians^11–13^. Music consists of many elements such as pitch, timbre, rhythm, intensity, melody, and harmony, all studied extensively in music psychology. For instance, pitch relates to the frequency of sound waves, while rhythm involves timing and patterns^14^. Melody and harmony shape musical dynamics, with concepts like contour linking the two^14^. Due to its complexity, music practice demands attention and repetition, which fosters brain plasticity^15–18^. Musicians, as a result, develop enhanced auditory sensitivity^19,20^ and a more efficient bilateral auditory cortex (AC)^21^. Furthermore, musical training has been shown to improve working memory (WM) in elderly individuals^22^.

Most studies on memory processes have focused on visuospatial stimuli, with auditory research primarily examining verbal sounds^1^. The initial memory process mainly studied with letters or numbers is sensory memory (SM), which serves as a temporary store for sensory information, including auditory stimuli, before it decays, is interfered with, or transitions to more durable storage. Research by Cowan^23^ and Deutsch^24^ has highlighted the limitations of auditory SM, revealing that individuals can only discern differences between similar sounds within a narrow temporal window. A memory process integrating a few items of information over minutes is WM, pivotal for complex cognitive tasks and reasoning^25^. In music, WM aids in recognizing and integrating sound attributes necessary for understanding and enjoying music. Lastly, long-term memory (LTM), the most durable memory system, is essential for recognizing familiar melodies and other auditory stimuli^26^.

The recent framework of predictive processing expands the understanding of memory, including auditory memory, as a dynamic process shaped by real-time listening and prediction^27,28^. Predictive coding (PC) involves a bidirectional flow of information within a hierarchical neural network^29^. In auditory perception, it describes how sound sequences are processed and compared to existing models, which are updated when sounds do not match these models^30^. Higher-level units send predictions to lower-level units, and when top-down predictions align with bottom-up sensory input, neural responses are suppressed, reflecting passive sensory adaptation. This predictive function engages nearly the entire brain^31^, with the cerebellum identified as a crucial area for higher-order perceptual and cognitive processing^28^. The cerebellum appears more involved in transforming memories and continuously updating semantic models used for event prediction rather than merely reactivating memories^28,32^.

To date, despite empirical evidence, the psychological and neural mechanisms underlying the acquisition, recognition, and retention of complex musical sounds remain scattered and unsystematic^1^. In turn, it is now well-established that uncorrected hypoacusia is a significant risk factor for cognitive decline and dementia^33^. Therefore, understanding the neural mechanisms for processing and retaining non-verbal sounds, particularly in aging populations, is crucial.

This systematic review aims to: (i) organize existing neuroimaging literature to identify the main brain mechanisms related to SM, short-term memory (STM), WM, and LTM for non-verbal auditory information; and (ii) explore variations in these neural memory processes based on participant characteristics, focusing on differences between musicians and non-musicians, younger and older individuals, and those with congenital amusia versus those without. Additionally, we conducted a coordinate-based meta-analysis to identify the brain structures most frequently associated with memory processes for non-verbal sounds.

## 2. Methods

### 2.1 Search strategy and study eligibility

This systematic review adhered to the guidelines outlined in the Cochrane Handbook for Systematic Reviews ^34^ and the Centre for Reviews and Dissemination ^35^. The review protocol was registered with the International Prospective Register of Systematic Reviews (PROSPERO: CRD 42022371132) before the beginning of the data extraction. The PRISMA statement ^36^ was followed in implementing the systematic approach. Comprehensive searches in electronic databases Scopus and PsycInfo were conducted, using relevant keywords and MeSH terms adapted to each database: “sound”, “auditory”, “tonal”, “music”, “pitch”, “timber”, “timbral”, “memory”, “learning”, “amusia”, “brain”, “cerebral cortex”, “lesion”, “clinical”, “behavioral”, “functional connectivity”, “structural connectivity” and not “verbal”, “speech” or “animal. Additionally, the reference lists of the included articles were manually examined for further relevant publications. The search results were imported into Rayyan ^37^ in RIS or XML format for screening.

Studies were included if they met the following criteria: 1) focus on non-verbal memory for sounds; 2) inclusion of adult participants aged 18 years and above; 3) inclusion of brain data obtained through neuroimaging methods. The exclusion criteria were as follows: 1) non-peer-reviewed studies; 2) studies published in languages other than English; 3) review articles; 4) meta-analyses; 5) animal studies; 6) post-mortem studies; 7) single case reports; 8) computational studies and 9) studies that focus solely on the mismatch negativity (MMN) response since it indexes automatic perceptual predictive processes rather than conscious discrimination memory processes ^38^.

Following the literature search, two researchers independently screened the studies based on titles and abstracts. Discrepancies were discussed with a third researcher. Subsequently, the full texts of the selected studies were retrieved and evaluated. A table was then created to extract the most pertinent data and to evaluate the methodological quality of the studies included in the review.

### 2.2 Data extraction

Each study was assessed for the following variables: general information such as author, article title, year of publication, journal, and geographical location; study characteristics including memory domain and study exclusion criteria; participant characteristics such as sample size, mean age, gender, clinical population, and musical expertise; and study methods including paradigm, measurement tools, type of stimulation, design, behavioral and brain outcomes, and analysis performed.

### 2.3 Quality assessment

Assessing the quality of studies across different research designs is a complex task as there is currently no universally recognized and scientifically proven tool accessible for this purpose. To comprehensively evaluate the quality of the studies included in this systematic review, the QualSyst tool ^39^ was utilized, providing a systematic and reproducible approach to quantitatively assess the quality of research across various study designs. As no qualitative studies were included in this review, only the checklist for assessing the quality of quantitative studies was utilized. For the quantitative studies, 14 criteria items were evaluated based on the degree to which they were met ("yes" = 2, "partial" = 1, "no" = 0). Items that were not applicable to a particular study design were marked as “not applicable” (NA) and were not included in the calculation of the summary score. Furthermore, since the review exclusively included observational studies and excluded interventional studies, items 5, 6, and 7 were deemed NA and were not scored. A summary score was then calculated for each paper by dividing the total score obtained by the total possible score. A score greater than 0.75 indicates high quality, between 0.55 and 0.75 indicates moderate quality, and less than 0.55 indicates low quality in the study. Quality assessment can be found in **Supplementary Table 2**.

### 2.4 Meta-analysis

We conducted an activation likelihood estimation (ALE) meta-analysis using GingerALE 3.0.2^40–42^, a software designed to assess the convergence of coordinates from independently conducted MRI studies examining the same construct. In this meta-analysis we compared the experimental conditions involving learning and memory processes (L) versus the control conditions non involving learning or memory processes (NL) at cluster level inference p < 0.01 (FWE).

Out of the 90 neuroimaging and neurophysiological studies reviewed, 34 studies reporting results in stereotactic coordinates (foci) within the Montreal Neurological Institute (MNI) or Talairach three-dimensional coordinate systems were identified. Among these, 15 studies that specifically compared L to NL were included in the quantitative synthesis. When coordinates were in Talairach space, they were converted to MNI space with the ‘tal2mni’ Matlab function.

## 3. Results

### 3.1 Characteristics of included studies

The PRISMA flowchart (**Figure 1**) for systematic reviews^36^ offers a comprehensive overview of the process of identifying, screening, assessing eligibility, and including studies. A total of 1,910 records were initially identified through manual and database searches. After removing duplicates, 1,729 articles were screened based on their title and abstract. Subsequently, 109 records underwent full-text review. Finally, 90 publications were included in the qualitative synthesis. Study characteristics can be found on **Supplementary Table 1**.

**Figure 1.**
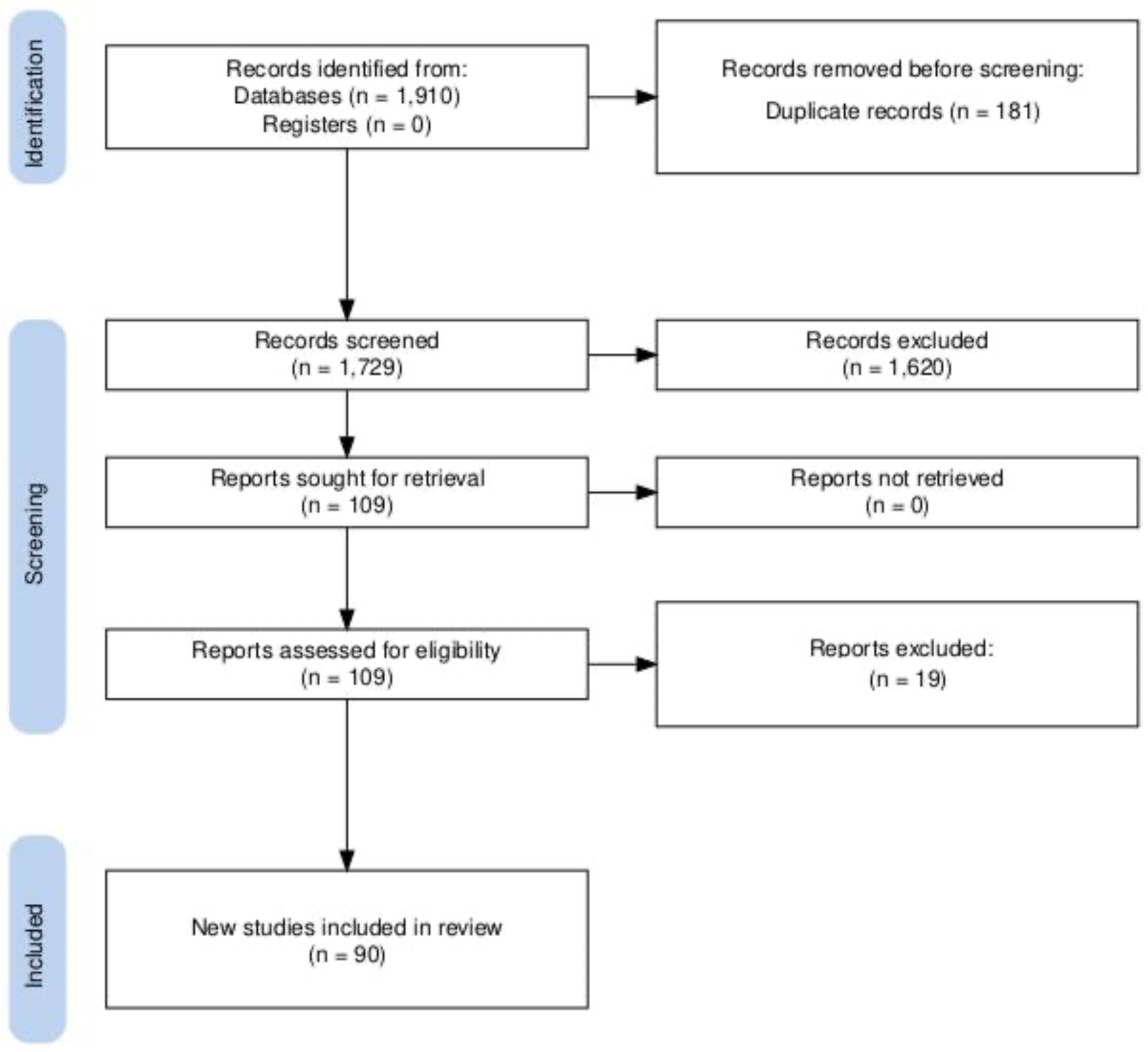
PRISMA flow chart illustrating the process of assessing the eligibility of neuroimaging studies included in the systematic review.

#### 3.1.1. Publication year and study location

The range of years for the studies included in the review is from 1990 to 2023, with a peak of 9 studies published in 2021 (**Figure 2**; **Supplementary Table 1)**.

**Figure 2.**
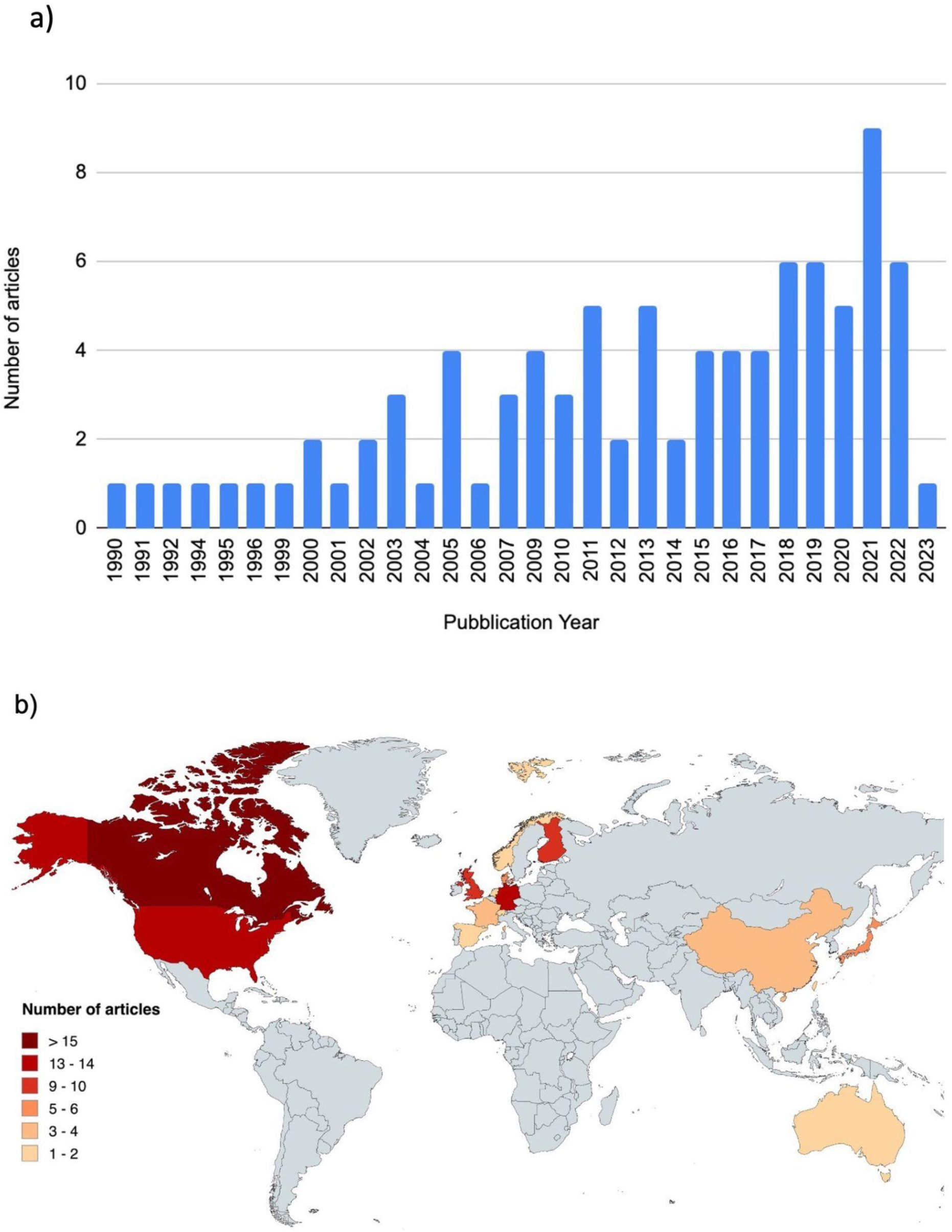
Number of articles (n=90) per publication year (a) and study location (b).

The geographical distribution of the studies is: Canada (16), Germany (14), U.S.A. (13), Finland (10), U.K. (9), Japan (6), Denmark (5), France (4), China (3), Netherlands (2), Switzerland (1), Spain (1), Norway (1), Taiwan (1), Australia (1), and not specified (3). This information emphasizes that research on the neural mechanisms underlying the storage of auditory memory information has been carried out mainly among Western listeners (**Figure 2**).

#### 3.1.2. Sociodemographic variables

The mean sample size of the enrolled subjects was 28.89 (interquartile range: 17.75), with the mean sample size of the included subjects at 27.85 (interquartile range: 16). Most studies (81; 90%) involved healthy participants, while nine studies (10%) focused on clinical studies. Among the clinical population, four studies (4.4%) involved patients with amusia, one (1.1%) involved patients with epilepsy, one (1.1%) included patients with Alzheimer’s Disease (AD) and Semantic Dementia, one (1.1%) involved AD patients with Posterior Cortical Atrophy (PCA), one (1.1%) involved patients with schizophrenia, and one (1.1%) included individuals with Mild Cognitive Impairment (MCI) and early AD.

Regarding gender, four studies (4.4%) did not specify the gender of participants^43–46^, while the remaining studies (95.5%) reported a female participant percentage of 52.46%. Age is recognized as a determinant factor influencing memory processes^47^; however, five studies (5.5%) did not report participant ages^43,48–51^. Eighteen studies (20%) provided only age ranges. In the remaining studies (74.5%), the mean age was 30.03 (interquartile range: 6.42), with ages ranging from 20 to 72. Six studies specifically focused on memory in aging.

Musical expertise was another variable examined, with 13 studies (14.44%) comparing musicians (with at least three years of training) to non-musicians. Among the subjects analyzed, 235 (53%) were musicians, while 208 (47%) were non-musicians.

#### 3.1.3. Exclusion criteria

The majority of studies (91.1%, N=82) clearly defined their inclusion criteria, which varied based on research objectives. The most common exclusion criteria included left-handedness, hearing impairment, and a history of neurological or psychiatric conditions. Specifically, 14 studies (17.07%) excluded participants with both hearing impairment and neurological/psychiatric disorders, while three studies (3.66%) excluded those with only neurological/psychiatric disorders. Additionally, seven studies (8.54%) excluded individuals with hearing impairment, five studies (5.49%) excluded left-handed participants, and four studies (4.88%) excluded those with both left-handedness and neurological/psychiatric disorders. Furthermore, 11 studies (13.41%) excluded participants with both left-handedness and hearing impairment, and 14 studies (17.07%) excluded participants with all three conditions.

Two studies excluded participants with absolute pitch—one in combination with hearing impairment^52^ and the other alongside left-handedness and neurological or audiological disorders^53^. Some studies also used musical expertise as an exclusion criterion, with 2.44% excluding non-musicians^54,55^ and various combinations of hearing impairment, left-handedness, and neurological/psychiatric disorders among musicians. One study excluded musicians without additional exclusion (1.22%)^43^, while four studies included musicians with hearing impairment (4.88%)^56–59^, one study included musicians with left-handedness (1.22%)^46^, one study included musicians with left-handedness and a history of neurological or psychiatric disorders (1.22%)^50^, and one study included musicians with hearing impairment and a history of neurological/psychiatric disorders (1.22%)^60^.

Other exclusion criteria encompassed medication usage, such as participants with left-handedness, a history of neurological/psychiatric disorders, hearing impairment, visual impairment, and current medication use (2.44%)^61,62^; those with left-handedness, medication use, and alpha dominant resting alpha activity (1.22%)^63^; participants with a history of alcohol or drug abuse, hearing impairment, a history of neurological/psychiatric disorders, and current medication use (1.22%)^64^; and those with hearing impairment, mental retardation, current medication use, and other neurological/psychiatric disorders aside from schizophrenia (1.22%)^65^.

The remaining studies exhibited diverse exclusion criteria. These included excluding participants with neurological or psychiatric issues and a history of chemotherapy (1.22%)^66^, those with hearing impairment and prior experience in psychoacoustic tasks or musical training (2.44%)^67,68^, participants with vision impairment and poor task performance (1.22%)^69^, individuals under 50 years old with mild hearing loss, recent medication changes, and neurological, psychiatric, or health problems (1.22%)^70^, as well as those with cerebrovascular disease, a history of hearing loss, or congenital amusia (1.22%)^71^.

### 3.2. Memory subsystems

For the purpose of systematic review, we have tried to distinguish studies by memory subsystems (SM/STM, WM, LTM), although there is often an overlap between them as measured by the various paradigms. If we consider memory subsystems, the largest proportion of studies focuses on WM (n=33), making up 36.6% of the total. This is followed by LTM (n=25), which accounts for 27.8%. Then we have SM and STM (n=17), which together account for 18.9%. After that we have studies that combine two memory subsystems: WM and LTM (n=7; 7.8%), SM and WM (n=6; 6.7%) and SM and LTM (n=2; 2.22%).

### 3.3. Methods of studies

#### 3.3.1. Measurement tools and paradigms

For SM and STM, the most commonly used neuroimaging tool is EEG (n=12). Other tools included fMRI (n=2), MRI (n=1), PET+MRI (n=1), and MEEG (n=1). The paradigms used in these studies include: “same/different” (n=2), discrimination tasks (n=1 for frequency, n=7 for pitch; n=1 sequences regularity), oddball (n=1), old/new paradigm (1), active and/or passive listening (n=2), rare tones perception task (n=1) probe tone task (n=1).

For WM, the most frequently used neuroimaging tool is fMRI (n=13), followed by EEG (n=12), MEG (n=4), MEEG (n=2), PET (n=1), and MEG + fMRI (n=1). The most used paradigms include: delayed match-to-sample task (n=9), same/different (n=7), N-back (n=4), discrimination tasks (n=1 for pitch, n=1 for frequency, n=1 for length, n=1 for sequences), oddball (n=3), digit span (n=2), segmentation task (n=1), two-interval two-alternative forced-choice paradigm (n=1), passive listening + WM task (n=1), melody listening + deviant sound recognition task (n=1).

For LTM, the most common tool is fMRI (n=9), followed by EEG (n=6) and MEG (n=4). Other tools include: MRI + MEG (n=1), EEG + Pupillometry (n=1), ECoG (n=1), fMRI + Polysomnography (n=1), and PET (n=1). The paradigms include: immediate and/or delayed recall (n=5), familiarity task (n=3), same/different (n=3), old/new (n=2), discrimination task (n=2), music or sounds listening (n=2), active learning (n=2), dual-phase paradigm (n=1), imaginary recall (n=2), oddball + statistical learning (n=1), melodies learning + deviants recognition task (n=1), and sound-to-picture matching task + familiarity tasks (n=1).

Six studies examined both SM and WM, with the majority using fMRI (n=5) and one using EEG. The most used paradigms include: pitch discrimination task + n-back (n=2), n-back + passive listening (n=2), frequency discrimination task (n=1) and deviant sound discrimination task (n=1).

Six studies investigated WM + LTM. Of them, 2 used the fMRI, 2 the EEG, 2 the combination of MEG+DTI and 1 the MRI+DTI. The paradigms include: old/new (n=2), continuous recall task (n=1), oddball (n=1), active listening + recall (n=1), rhythm and melody learning tasks (n=1), and probabilistic reinforcement learning task (n=1).

Lastly, two studies investigated both SM and LTM, both using fMRI. One study combined fMRI with skin conductance response, using paradigms like passive listening + recall and incidental sound detection task + encoding.

EEG is the most frequently used neuroimaging tool, accounting for 39.44% of the studies, immediately followed by fMRI (38.33%), MEG (11.11%), MRI (5%), PET (2.77%), and ECoG (1.11%).

#### 3.3.2. Auditory stimuli

The auditory stimuli employed are diverse and functional to the design of specific studies. They can be broadly categorized into two groups: studies that utilized tones or sounds (67.78%) and studies that instead employed melodies or songs (32.22%).

#### 3.3.3. Design

Most of the included studies are cross-sectional/observational type (87.78%). Ten studies (11.11%) are case-control studies, while only one study (1.11%) is a cross-cultural study.

### 3.4. Quality assessment

The quality of the studies included in this review was evaluated comprehensively using the QualSyst tool, in accordance with the guidelines provided by Kmet et al.^39^. The total scores for each study ranged from 0.59 to 1.0 with a mean of 0.93 ± 0.10. In general, the studies examined showed good practices with high scores in items related to object and design description, as well as the interpretation of results. However, they received lower scores in items related to subject characteristics and sample size (**Supplementary Table 2**).

### 3.5. ALE meta-analysis

The quantitative analysis of the main outcome included 15 studies, 1317 foci, 15 experiments, and a total of 439 participants. The contrast L > NL resulted in a single significant cluster encompassing the medial frontal gyrus (MFG), cingulate gyrus and superior frontal gyrus (SFG), mostly in the left hemisphere (**Figure 3**). The cluster analysis revealed a volume of 2.248 mm³, spanning from coordinates (-4, 0, 30) to (8, 18, 64), with a center at (1.1, 6.7, 47.2). The cluster contained 6 peaks, with the highest ALE value being 0.0271, corresponding to a p-value of 3.50E-8 and a Z-score of 5.39 at coordinates (2, 10, 40). The contrast NL > L did not yield any significant cluster.

**Figure 3.**
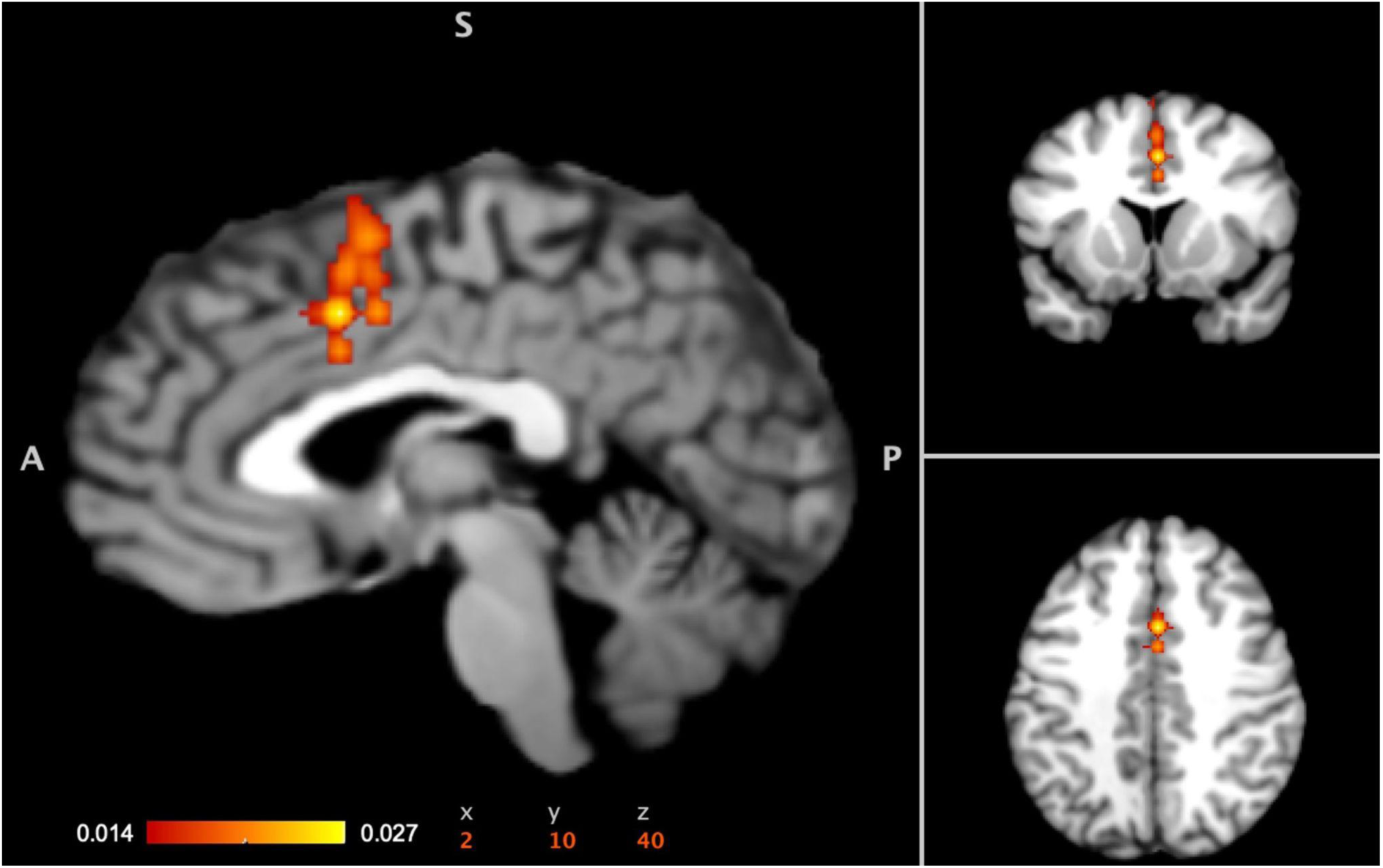
Anatomic estimation of the highest ALE value for the contrast L>NL overlapped with MNI template which revealed a significant cluster encompassing the medial frontal gyrus (60.6%), cingulate gyrus (38%), and superior frontal gyrus (1.4%), in both the left (91.5%) and right (8.5%) hemispheres.

## 4. Discussion

The objective of our systematic review and meta-analysis was twofold: to comprehensively organize the existing neuroimaging literature to identify the key brain mechanisms responsible of memorizing nonverbal auditory information, and to explore these neural memory processes across individual differences such as age, musical expertise and music-specific disorders (congenital amusia). An important finding of the systematic review is the difficulty in organizing the findings according to the memory subsystems as identified in the classical psychological literature. The reviewed studies seem to support a view of memory as a dynamic unified system for prediction of future events based on reconstruction from past memories. However, for coherence with the initial scopes of the review, we will discuss the findings according to the classical distinction in SM/STM, WM and LTM.

### 4.1. Memory domains

For the purposes of this review, we chose to group SM and STM together, as there is often a conceptual overlap between these two systems. For instance, Cowan^23^ and Massaro^72^ conceptualized two types of auditory SM: a perceptual store, which aligns with the classical SM system^73^, and a synthesized store, where sensory information is passively held for about 10 to 20 seconds and is partially analyzed, overlapping with the classic definition of STM^74^. Since then, many models have been proposed, with Schröger^75^ arguing that auditory SM cannot be regarded as a unitary phenomenon. Instead, it should be viewed as an overarching concept that encompasses auditory representation, integration, and short-term storage of information, thereby serving as a bridge that connects pre-representational auditory information and mature cognitive representations.

#### 4.1.1. Sensory and short-term memory in the brain

In the reviewed studies on discrimination memory for non-verbal sounds, primarily focusing on pitch and timbre features, a finding is the involvement of the right hemisphere, with a network involving the right temporal neocortex and the right frontal lobe responsible for maintaining the acoustic information needed to perform discrimination tasks^63^.

Another important aspect is the effect of adaptation and habituation to repeated stimuli on neural responses to sounds^76–79^. According to Stein’s classic theory^80^, a novel stimulus initially activates the excitatory mechanism, followed by an inhibitory response. With repeated exposure, the inhibitory mechanism becomes conditioned to the stimulus onset. As the stimulus becomes more predictable, this conditioned inhibitory response begins to override the direct excitatory response, resulting in diminished neural responses to the repeated stimulus^80^. Two studies utilizing the N1 evoked response to frequency and beat changes demonstrate that repetition leads to habituation of neuronal activity, indicated by decreased neurophysiological responses to the repeated sound features, depending on their similarity to the initial features^81,82^. These findings suggest that habituation may serve as a fundamental mechanism of sound memory over brief time spans, further supported by results from naturalistic listening paradigms^83^.

Additionally, functional connectivity (FC) within the auditory network, specifically between the superior temporal gyrus (STG) and the inferior parietal lobule (IPL), appears to be dynamically modulated depending on the characteristics of the auditory STM tasks (e.g., pitch discrimination or n-back tasks), and also when sounds are presented during a visual task^84^. This challenges the notion of a hierarchically organized AC^85–87^, suggesting that information flow in the human AC is more heterarchical. IPL also seems to have a significant role in analyzing both task-irrelevant and task-relevant auditory information^84^.

Lastly, the amplitude of the sustained anterior negativity (SAN) seems to increase with the number of STM retained items, supporting the hypothesis that the SAN is an electrophysiological index of brain activity related to the maintenance of auditory information in STM^88^.

#### 4.1.2. Working memory

The examined studies identified a widespread network of brain areas involved in auditory WM within both the cerebrum and cerebellum, crucial for the predictive mechanisms of memory. The hippocampus is central to this network, showing significant activity and connectivity with the AC and frontal structures, aligning with its established role in memory processes across various sensory domains^1,28,89,90^. Other key regions include the hippocampus and parahippocampal areas, particularly active during the encoding of complex auditory scenes and sequences. Increased activation with memory load was found in the anterior hippocampus, angular/supramarginal gyrus (SMG), and the secondary AC. Additionally, the superior parietal cortex seems to be involved in both visual and acoustic STM^43^.

Factors like motif repetition influence hippocampal activity and its connections with other WM-related areas, such as the dorsolateral prefrontal cortex, supplementary motor area, and cerebellum, indicating its role in encoding musical motifs and supporting LTM formation^54^. The neural connections between the AC, hippocampus, and inferior frontal gyrus (IFG) supports a system for auditory WM that maintains sound-specific representations through projections from higher-order areas^56^. Interestingly, this network appears to be nonspecific.

Additional key areas include the auditory cortices, specifically Heschl’s gyrus (HG) and the planum temporale (PT). Research indicates that the discriminability of conditioned tones is heightened in the AC, particularly in HG, during both reinforced and non-reinforced contexts^91^. HG is well-known for its role in pitch discrimination; deviant patterns trigger increased neural activity and left-lateralized connectivity from HG to PT, as well as heightened excitation within left HG^57^. For instance, fMRI imaging revealed that strong-learners exhibited increased activity in the left HG, left posterior superior temporal gyrus, and SMG following training, while weak learners displayed heightened activity in the left HG and anterior insula (AI), alongside involvement of lingual, orbitofrontal, and parahippocampal regions. The SMG is implicated in auditory memory storage, differentiating performance levels in pitch memory tasks^92^. Moreover, activation in the left PT positively correlates with n-back task performance^93^.

Other frontal structures, particularly the IFG and ACC, are also involved in auditory WM, reflecting attentional and cognitive control processes. Functional dissociation within this network suggests distinct regions handle spatial versus non-spatial auditory features or rhythm versus melody^94^, with the IFG specifically involved in distinguishing tones held in memory^56^. Decreased activation in the right IPL and superior right middle frontal gyrus occurs during less demanding tasks, indicating reduced attentional control demands^95^. The ACC is associated with effortful listening, evidenced by increased neural oscillation power in the theta frequency band (around 4-8 Hz) over frontal midline scalp regions, as well as in posterior alpha during challenging retention intervals^96,97^. This aligns with the idea that alpha oscillations (around 8-12 Hz) may serve as a mechanism for disengaging task-irrelevant regions^50^.

The cerebellum, especially lobules VII-VII, along with parietal and prefrontal structures, is engaged in WM tasks with increased cognitive load^98^ and is activated during rhythm WM, along with the vermis, right AI, and left ACC^99^.

EEG and MEG studies highlight modulation of memory-related brain activity by sound and person-related features. Anurova et al.^100^ identified distinct networks generating P300 and the positive slow wave during auditory n-back tasks, supporting the dual-stream model for auditory processing. Indeed, location and pitch WM tasks generate slow potentials influenced by information volume and reduced by distractors^101,102^. Early components of auditory evoked responses differ between spatial and non-spatial WM tasks, suggesting separate neuronal networks for initial sound identification and WM encoding^103^.

Furthermore, auditory spatial tasks are associated with higher gamma-band spectral amplitude (approximately 50-65 Hz and 90-100 Hz) at right posterior and central sensors. This increased activity lasts several hundred milliseconds after the task cue offset, highlighting distinct cortical generators for encoding versus delay-related activations. Additionally, no task-related modulation occurs below 40 Hz, suggesting that alpha-band activity is not involved in task set selection^104^.

Finally, even emotion-related features of sounds modulate WM mechanisms, with the right amygdala and left precuneus showing task-related decreases in activity influenced by the emotional connotation of the stimulus^105^.

#### 4.1.3. Long term memory

Many studies focused on LTM. For instance, Czoschke et al.^69^ studied the neural representation of pitch and location for memorized sounds, revealing feature-selective activity for both features in the auditory cortex, superior parietal lobule, and SMG. Notably, no regions exclusively encoded pitch memory. Their findings suggest a hierarchical organization within the auditory system, with superior decoding accuracy for location compared to pitch, indicating the brain’s ability to integrate features within the same region^69^.

Daikhin and Ahissar^106^ investigated the neural mechanisms of frequency discrimination, showing that cognitive strategies shift based on task conditions. In a "no-reference" condition, increased activation occurred in frontoparietal areas associated with WM, whereas the presence of a reference tone facilitated faster learning and reduced activity in areas linked to stimulus retention. This transition from WM reliance to LTM highlights the brain’s adaptive capacity in optimizing information processing.

Other research on auditory memory retrieval after musical exposure revealed distinct cortical activity patterns during listening and retrieval. Ding et al.^107^ observed differences in high-gamma band activity initiation during music listening, primarily originating from the temporal lobe and SMG, while recall showed initial activity in the IFG and precentral gyrus. Additionally, delta and high-gamma band responses in the SMG, temporal, and frontal lobes correlated with music intensity fluctuations, exhibiting noticeable temporal delays. Frontal lobe activity lagged behind temporal activity during listening but preceded it during recall, indicating the involvement of both bottom-up and top-down processes in music processing.

In a series of MEG studies^108,109^, Brattico’s team adapted the old-new paradigm, typically used for visual recognition memory, to investigate non-verbal auditory encoding and recognition memory in a naturalistic setting. They found that simpler tonal sequences activated hippocampal and cingulate regions, while more complex atonal sequences primarily engaged the auditory processing network, indicating that neural pathways change qualitatively based on stimulus complexity^108^. Further analysis^109^ revealed that successful recognition of previously learned musical pieces involved a broad brain network, including the cingulate gyrus, hippocampus, insula, inferior temporal cortex, frontal operculum, and orbitofrontal cortex. Additionally, positive correlations were observed between auditory sequence recognition and WM abilities, particularly in the medial cingulate gyrus and regions not typically associated with auditory processing, such as the inferior temporal, temporal-fusiform, and postcentral gyri, suggesting an interplay between different memory subsystems.

The role of the insula was also highlighted in a study by Mutschler et al.^110^, who used fMRI under active and passive learning conditions. Their findings indicated differential brain responses to action-related sounds in the insula, particularly in the hand sensorimotor area, suggesting the insular cortex’s involvement in the initial phase of learning action-perception associations. Additionally, Zioga et al.^111^ conducted a study where participants learned melodies based on an artificial musical grammar and then composed new melodies. EEG recordings during melody judgment showed that participants successfully acquired the music grammar: delta band power during initial exposure correlated positively with learning, while post-training alpha and beta band power indicated neural retrieval mechanisms. Learning emerged as a significant predictor of creativity, as evidenced by the P200 response to incorrect notes, indicating a connection between the neural foundations of learning and creativity^111^.

Tseng et al.^112^ investigated the timeline of brain dynamics during music memory retrieval, revealing that low-frequency oscillations in the frontal, temporal, and parietal regions increased during the processing of learned melodies and distractors. Cross-frequency coupling between low-frequency phases and high-frequency amplitudes intensified in the frontal and parietal regions when participants successfully distinguished between targets and distractors. Theta and alpha band phase locking was also observed during music memory retrieval, indicating the presence of FC and phase-amplitude coupling in the neocortex^112^.

Anodal transcranial direct current stimulation (tDCS) provides evidence for the causal involvement of the AC in auditory memory processes. For instance, tDCS applied to the right AC disrupted pitch discrimination learning for up to two days^113^, while stimulation of the left AC or sham stimulation had no effect. This disruption correlated with a reduction in the N1m component amplitude in the right AC, measured by MEG, indicating a causal relationship between right AC responses and fine pitch processing^113^.

Some studies explore learning using sound stimuli or short melodies instead of full musical compositions. For example, Vaquero et al.^60^ discovered that the volume of the right arcuate fasciculus, particularly in its anterior segment, can predict the rate and speed of learning in rhythm tasks. Additionally, fractional anisotropy in the right long segment can predict the rate of melody learning, supporting the involvement of the arcuate fasciculus in auditory-motor learning and the presence of feedback-feedforward loops during auditory-motor integration.

The right hippocampus is another critical area for LTM of sound stimuli, exhibiting greater responsiveness during recognition and retrieval than the left hippocampus^114^. Moreover, the left IFG shows stronger activation than the right IFG in response to sound stimuli^114^. The hippocampus is activated not only during recognition or retrieval^109,115^, but also during the implicit learning of acoustic sequences, highlighting its role in memory formation for perceptual-based representations, regardless of whether the learning is implicit or explicit^116^.

Another approach to studying recognition memory involves contrasting auditory stimuli, typically music, based on self-assessed familiarity. For example, Jagiello et al.^117^ found that listening to familiar musical pieces led to significant pupil dilation from 100 to 300 ms after stimulus onset, indicating rapid activation of the autonomic salience network. An EEG study also revealed brain responses similar to those observed in classic old/new memory retrieval paradigms when contrasting familiar with unfamiliar songs^117^.

### 4.2. ALE meta-analysis

Our meta-analysis revealed a significant cluster in the MFG extending to the cingulate gyrus and SFG, predominantly in the left hemisphere. The cingulate gyrus, involved in episodic memory processing^118^, plays a fundamental role in auditory memory encoding^119^ and recognition^108,109,120^. Moreover, it is also associated with context-related priming, along with the medial and dorsal superior frontal gyri, midline structures, and right angular gyrus^121^. Additionally, the cingulate gyrus is involved in memory recognition in WM and LTM tasks, together with the SFG^122^.

The SFG, MFG, and cingulate gyrus are part of the posterior default mode network (DMN). Bar^123^ suggested an overlap between the medial prefrontal cortex (mPFC) and the posterior cingulate cortex (PCC), as well as areas involved in associating memory stimuli with prior experiences^28^. The SFG has been shown to modulate theta and alpha frequency bands during encoding, coordinating memory processes through low-frequency oscillations^124^. Additionally, DMN regions are activated during the listening of emotional music^125^. For instance, the medial, superior, and middle frontal gyri, along with the cingulate gyrus, are engaged when participants evaluate the emotional expression of music^126^. In another study, musical pieces with positive personal valence compared to less positive pieces activated the left MFG, left precuneus, right SFG, left PCC, bilateral middle temporal gyri, and left thalamus^127^.

### 4.3. The role of individual differences

We found many studies related to how individual differences, such as gender, age and musical expertise, may influence how the brain processes non-verbal sounds and music. For instance, gender differences were observed in the activation patterns of both anterior and posterior perisylvian regions using fMRI during a pitch memory task^128^. Male subjects showed stronger left-hemisphere activation than the right during both "perceptual" and "memory" phases. Males also had more cerebellar activation, while females exhibited greater PCC/retrosplenial cortex activity. Despite these brain differences, behavioral performance was similar, indicating males rely more on left-hemisphere processing for basic pitch tasks, similar to patterns seen in language studies^128^. P300 studies show that age, musical training, and cognitive abilities influence neural signatures of auditory WM, with males showing larger P3b amplitudes and females showing greater differentiation between targets and distractors in auditory oddball tasks^62^.

In the next paragraph we will consider the role of individual differences in terms of musical expertise, age and congenital amusia, a deficit in melody perception and production that cannot be explained by hearing loss or any type of brain damage.

#### 4.3.1. Musical expertise

Numerous studies comparing musicians with non-musicians, revealed that musicians tend to outperform non-musicians in visual, phonological, and executive memory tasks. Musicians exhibit faster WM updates in both auditory and visual domains, indicated by shorter latency P300s and larger P300 amplitudes in response to auditory stimuli^61^. Additionally, musicians exhibit larger blood-oxygen-level-dependent (BOLD) responses in brain regions associated with attention and cognitive control, particularly during challenging WM tasks^129^.

A study by Teki et al.^130^ on professional piano tuners found increased gray matter volume in several brain regions, including the right frontal operculum, right superior temporal lobe, anterior hippocampus, parahippocampal gyrus, and superior temporal gyrus, along with increased white matter in the posterior hippocampus. These changes correlated with tuning experience duration and starting age, highlighting the role of these regions in sound exploration and memory consolidation in an experience-dependent manner^130^. Furthermore, pianists demonstrate greater improvements in pitch and rhythm accuracy during auditory learning compared to motor learning, while non-musicians show greater rhythm improvements during auditory learning. Pianists’ motor responses during auditory learning correlate with pitch accuracy and auditory-premotor network activity, whereas non-musicians exhibit greater parietal responses linked to pitch accuracy^52^.

Differences between musicians and non-musicians are also reflected in EEG components. While repeated exposure to musical excerpts does not affect early neural mechanisms of music-syntactic processing (measured by early right anterior negativity, ERAN), it does modulate the amplitudes of the late positive component (LPC) and P3a^131^. Musicians exhibit enhanced plasticity in event-related potential (ERP) responses to deviant sounds, with activation in left and right temporal generators and a left frontal source, but not in the right frontal source, indicating its unique role in deviant sound processing^132^.

Mathias et al.^133^ explored how musical expertise influences auditory-motor associations in skilled pianists, finding that altered pitches elicited a larger N2 ERP component compared to original pitches, particularly in melodies learned through production rather than perception alone. This underscores the interplay between auditory perception and motor planning shaped by experience^133^. Lastly, musical training may enhance statistical learning of rhythm, with various types of training affecting neural representations of temporal statistical learning differently^53^. Overall, musical expertise seems to lead to extensive reorganization of processing pathways connecting superior temporal sources with the left IFG, which is involved in statistical learning^134^.

#### 4.3.2. Aging

Given the use of music as a factor promoting cognitive reserve and neurorehabilitation, some studies examine how aging affects the brain’s ability to memorize non-verbal sounds. Halpern et al.^135^ found that, analyzing expectations for musical sound, younger listeners relied more on specific melodic sequences, while older listeners focused on tonal patterns related to the musical key. Both groups had similar N1 responses; older adults showed stronger P200 and LPC activity, particularly in frontal regions, while younger adults’ LPC was more posterior. This suggests that older adults refine their musical understanding with age-related changes in neural activity supporting this ability^135^. Moreover, also P300 seems to be affected by age, with latency decreasing and amplitude increasing with aging^45^.

Similarly, Koelsch et al.^136^ investigated neural activations when listening to irregular chords in both adults and children. In adults, various brain regions were activated, including the IFG, orbital frontolateral cortex, AI, ventrolateral premotor cortex, STG, SMG, and superior temporal sulcus. These regions constitute different networks involved in cognitive and emotional aspects of music processing. In children, the activation pattern in the right hemisphere was similar to that of adults, but in the left hemisphere, adults exhibited larger activations in the prefrontal areas, SMG, and temporal areas. Both musical-trained adults and children showed enhanced activations in the frontal operculum and the anterior segment of the STG. Similarly, Sikka et al.^46^ found greater activation in the left STG for younger adults, and in the left superior frontal, left angular, and bilateral superior parietal regions for older adults during melody recognition^46^.

Thaut et al.^137^ showed that exposure to familiar music in elderly activates brain regions associated with autobiographical memory and emotional responses, including the mPFC, precuneus, AI, basal ganglia, amygdala, hippocampus, and cerebellum. Given that these areas are minimally affected by early-stage AD, except for the hippocampus^119^, this bilateral network of prefrontal, emotional, motor, auditory, and subcortical regions may provide insights into the preservation of musical LTM in cognitively impaired older individuals. Increased FC between auditory regions and the mPFC, in combination with reward and DMN areas when listening to familiar and pleasant music, was also observed^70^, supporting the idea that music listening can offer a mechanism for music-based interventions, promoting healthy aging by providing an auditory channel towards the mPFC.

Finally, Hsieh et al.^138^ found that AD patients had mild deficits in auditory semantics, with recognition of famous tunes linked to atrophy in the right anterior temporal lobe, especially the temporal pole. This area, critical for face identification^139^, was separated from regions involved in everyday sound recognition. Additionally, patients with semantic dementia who retained musical knowledge showed less atrophy in the right temporal pole. The study suggests that the right temporal pole is vital for processing known tunes and faces, reflecting a common recognition mechanism for unique identities^138^. Moreover, AD patients seem to have reduced activation in the right IFG for musical semantic memory, while for incidental musical episodic memory, an abnormally enhanced activation was localized to the precuneus and PCC in both AD patients and those with PCA^71^.

#### 4.3.3. Amusia

Studies by Peretz’s team^58,59,140^ focused on investigating music-specific processing and memory deficits in individuals with congenital amusia, a disorder characterized by “difficulty in perceiving, or making sense of, music”^141^. The most common form of this disorder involves impairments in detecting, recognizing, and retaining tunes in memory^142^. Evidence indicates that neural responses, such as MMN and N1, are preserved in the automatic and early stages of encoding music-relevant sound features like pitch. However, deficits emerge in conscious memory processes, as shown by abnormal brain responses, including larger N2 and P3 components compared to controls. Notably, the P3b response for larger pitch differences remains normal, indicating that the amusic brain can detect quarter-tone pitch differences, suggesting intact neural circuitry for fine-grained pitch perception at a preattentive level. However, there are deficits in the conscious recognition of these distinctions in pathways beyond the AC. Consequently, it has been proposed that the AC’s representation of pitch in amusia is not effectively linked to musical pitch-related knowledge via the auditory-frontal neural pathway^140^.

### 4.4. Music and language

Recently, there has been a surge of interest in the intersecting neural networks responsible for both music and language processing. Music and language domains both comprise hierarchically structured sequences which seem to demand common syntactic processing in the brain^143^. The studies included in the present review suggest activations in the areas of the brain related to linguistic syntax processing while engaging in musical tasks. For instance, Sternin et al.^144^ used fMRI to explore how neural networks for music and language interact with memory, using different familiar and unfamiliar stimuli (*i.e.*, whole music, instrumental, acapella, only words). Results showed activations in the bilateral planum polare and left IFG across all conditions, with stronger bilateral planum polare activations in response to musical stimuli, supporting its role in both linguistic^145^ and musical^146^ syntax processing. Similarly, the left IFG, known for its role in both musical^147^ and linguistic^148^ syntax processing, showed greater activation for spoken than instrumental conditions, reflecting differing syntactic demands in music and language. Similarly, Koelsch et al.^149^ found overlapping neural activity for WM during the rehearsal and storage of both verbal (syllables) and tonal (pitches) information, suggesting shared resources for both processing. Quinci et al.^70^ further explored music-based interventions for healthy aging, revealing enhanced brain connectivity in both the basal ganglia and language networks after the intervention, linking musical familiarity to shared networks for memory of linguistic and musical sounds.

The Shared Syntactic Integration Hypothesis suggests that linguistic and musical information is processed, at least partially, through a common syntactic mechanism^150^. Evidence supports that WM plays a key role in this process by supporting the temporal storage and integration of both music and speech sounds. In an attempt to explore the topography of WM for music in naturalistic condition, Burunat et al.^54^ using fMRI, asked participants to perform a real-time music segmentation task, revealing activations in the opercular and triangular parts of the right IFG—homologous to Broca’s area, which is critical for linguistic syntax processing, and thus suggesting a WM involvement for syntax processing in two domains. In line with these findings, a MEG study by Maess et al.^147^ found that an ERAN was generated in response to out-of-key chords in Broca’s area and its right hemisphere homologue.

Another fMRI study suggestive of interaction between music and language through WM provides evidence in the context of statistical learning^151^. Researchers used an artificial grammar learning paradigm for memorizing musical patterns to examine neural mechanisms that process auditory stimuli with local dependencies (*i.e*. triplets of Shepard tones and percussion sounds). Standard triplets followed a simple statistical rule, whereas the deviant triplets violated this rule. Deviants’ recognition activated a broad frontostriatal and temporal network closely resembling the one involved in artificial grammar learning of speech sounds, suggesting shared WM mechanisms for statistical learning and prediction across both music and language domains.

The findings can also be interpreted through the frame of PC^30^, as it provides valuable insights to understand predictive processes in perception and action. As opposed to traditional models of perception, listening to music can be considered as an active state of action as it seems to offer epistemic stimuli for statistical learning. A key component of the PC framework is precision filtering, which involves generating another prediction about how precise the prediction is likely to be. This allows the brain to prioritize processing the more precise information rather than less precise one. Indeed, precision filtering requires maintenance of the relevant information in WM, which is where both linguistic and musical stimuli are stored and processed.

Another study by Bonetti et al.^152^ aimed to investigate the whole-brain FC involved in the rapid encoding of musical tones and its subsequent memory representation. Authors found that the network of brain areas detected in the study comprised areas mostly related to auditory, memory, attentional, and evaluative processes. Additionally, individuals with high memory skills showed a stronger activation in frontal operculum which is known for its critical contributions to phonological processing of speech sounds^153^. In line with these findings, Bonetti et al.^154^ reported activation in frontal operculum, along with other higher-level brain regions, in response to processing errors related to cognitive priors in music perception and cognition, especially in musicians. Similarly, another study by Yamashita et al.^155^ investigated the effect of lifelong musical instrument training on age-related cognitive decline and brain atrophy. Results revealed that musicians had superior verbal fluency compared to non-musicians, which can be attributed to the common phonological features in music-making and language production. In particular, considering that musical training facilitates memory and statistical learning about the acoustic structure of musical pieces, musical training may facilitate higher performance of letter fluency in language domain for musicians. Taken as a whole, these discoveries support the concept that PC and WM provide a suitable framework for interpreting language and musical processing.

## 5. Conclusions and limitations

A limitation of our paper is the significant methodological heterogeneity among the studies included, which, particularly for the meta-analysis, could introduce confounding factors that might affect the results. One of the challenges in writing this review was classifying the studies into the traditional categories of “SM”, “WM”, and “LTM”. Some studies, despite employing very similar methods and tools, were classified differently—even by the same authors.

Nonetheless, this review addresses a significant gap in the literature, where a substantial body of research on non-verbal memory processes has accumulated without any systematic overview. Only recently, a systematic review of findings on neurophysiological responses in aging has been completed^156^, although this is only marginally relevant to our memory-focused objectives. Additionally, a mini-review of the limited neuroimaging literature on musical working memory has been published^157^. Therefore, our review represents a significant step forward in the investigation of memory processes for non-verbal sounds.

## CRediT authorship contribution statement

**FFC:** Validation, Methodology, Formal analysis, Data Curation, Investigation, Writing – original draft, Writing – review & editing. **FC:** Investigation, Validation, Writing – review & editing. **BY:** Validation, Investigation, Writing - original draft (contribute to section 4.4), Writing – review & editing. **CG:** Validation, Writing – review & editing. **LQ:** Investigation, Writing – review & editing. **DR:** Writing - review & editing. **EB:** Conceptualization, Validation, Methodology, Supervision, Investigation, Resources, Writing – review & editing.

## Acknowledgements

Center for Music in the Brain (MIB) is funded by the Danish National Research Foundation (project number 117).

FFC Ph.D. Scholarship was funded by POR Puglia FESR FSE 2014-2020, CUP: H99121006630008.

This publication was also produced with the co-funding European Union - Next Generation EU, in the context of The National Recovery and Resilience Plan, Investment Partenariato Esteso PE8 "Conseguenze e sfide dell’invecchiamento", Project Age-It (Ageing Well in an Ageing Society), CUP: B83C22004800006.

